## Supplementary materials for "Targeting host deoxycytidine kinase mitigates *Staphylococcus aureus* abscess formation"

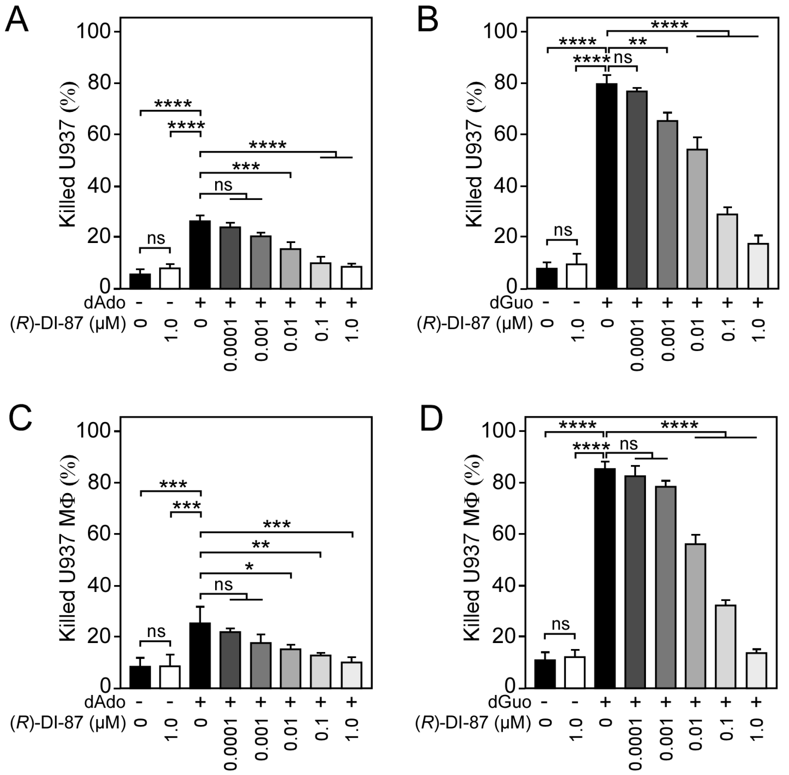


**Supplementary Figure 1. (*R*)-DI-87 prevents death-effector deoxyribonucleoside-triggered immune cell death in a dose-dependent manner. (A-D)** Survival rates of human U937 monocyte-like cells (U937) (A, B) or U937-derived macrophages (U937 MΦ) (C, D) exposed to dAdo or dGuo in the presence (+) or absence (-) of various concentrations of (*R*)-DI-87. Cells were also exposed to the inhibitor or vehicle only. 100 µM (A, B) or 200 µM (C, D) of dAdo or dGuo were used to treat the cells. Cell survival rates were analyzed 48 h post-treatment. Data are the mean (± standard deviation [SD]) values from three independent determinations. Statistically significant differences were analyzed with one-way analysis of variance (ANOVA) and Tukey's multiple-comparison test; ns, not significant ($P \geq 0.05$); *, $P < 0.05$; **, $P < 0.01$; ***, $P < 0.001$; ****, $P < 0.0001$.

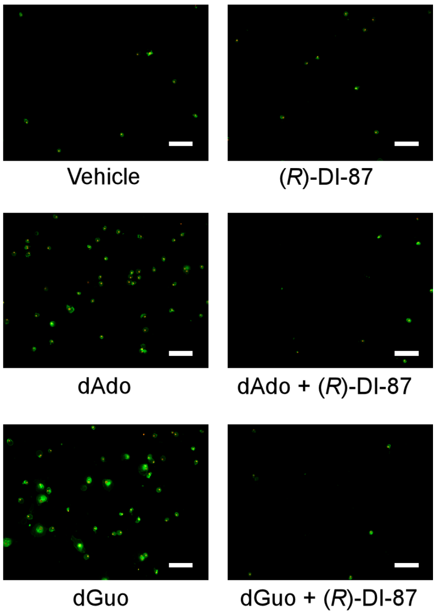


**Supplementary Figure 2. Inhibition of dCK prevents death-effector deoxyribonucleoside-mediated induction of immune cell apoptosis in primary human macrophages.** Analysis of (*R*)-DI-87-dependent prevention of host cell apoptosis via immunofluorescence microscopy. Primary human monocyte-derived macrophages (HMDMs) were exposed to dAdo or dGuo in the presence or absence of 1 µM (*R*)-DI-87 and stained using FITC-annexin-V/PI. Controls are indicated. White bars depict a length of 100 µm. Representative images are shown. 200 µM of dAdo or dGuo were used to treat the cells. Apoptosis rates were analyzed 24 h post-treatment.

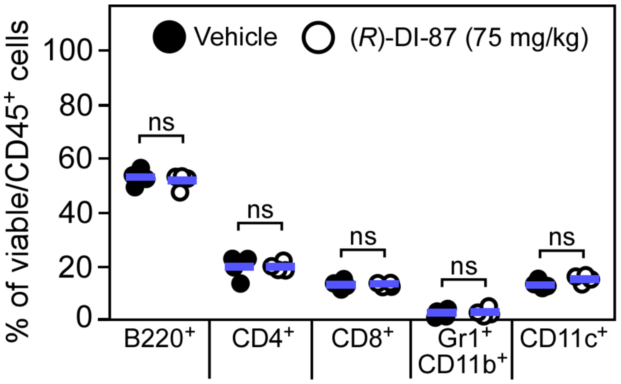


**Supplementary Figure 3. Immuno-phenotypic assessment of murine spleen tissues following continuous dCK inhibitor treatment.** Cohorts of female C57BL/6 mice were treated with (*R*)-DI-87 (75 mg/kg) or vehicle (40% Captisol) via oral gavage in 12-hour intervals for 23 days. On day 23, spleen tissues were collected and subjected to a FACS-based immuno-phenotyping approach. Statistically significant differences were analyzed by a two-tailed Student's t-test. ns, not significant ($P > 0.05$).

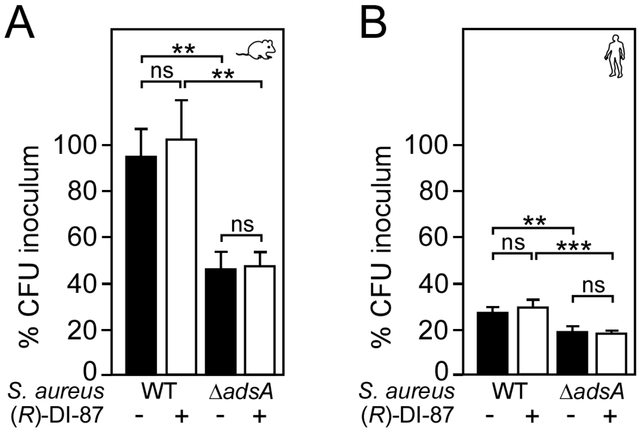


**Supplementary Figure 4. (*R*)-DI-87 does not interfere with staphylococcal survival in blood. (A, B)** Survival of wild-type *S. aureus* Newman (WT) or its *adsA* mutant (Δ*adsA*) in mouse (A) or human blood (B) in the presence (+) or absence (-) of (*R*)-DI-87 after 1 h of incubation. Data were recorded as percent inoculum. For experiments with human blood, three independent donors have been used. Statistically significant differences were analyzed by two-way analysis of variance (ANOVA) followed by Tukey's multiple-comparison test; ns, not significant ($P \geq 0.05$); **, $P < 0.01$; ***, $P < 0.001$.

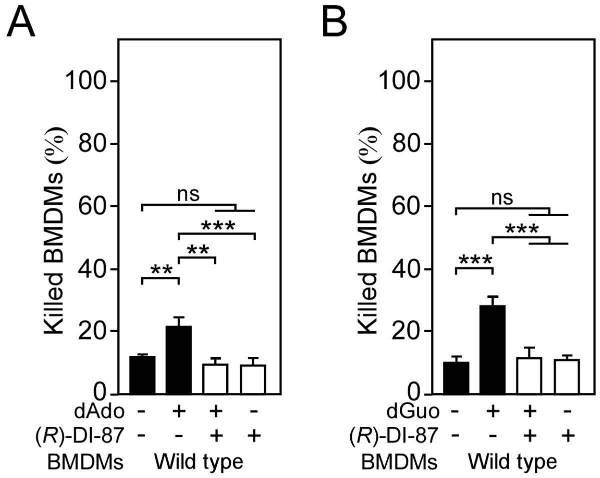


**Supplementary Figure 5. (*R*)-DI-87-mediated inhibition of dCK shields male mice-derived phagocytes from death-effector deoxyribonucleosides. (A, B)** Survival rates of male mice-derived bone marrow-derived macrophages (BMDMs) exposed to dAdo (A) or dGuo (B) in the presence (+) or absence (-) of 1 µM (*R*)-DI-87. Cells were also exposed to the inhibitor or vehicle only. 200 µM of dAdo or dGuo were used to treat the cells. Cell survival rates were analyzed 48 h post-treatment. Data are the mean (± standard deviation [SD]) values from three independent determinations. Statistically significant differences were analyzed by one-way analysis of variance (ANOVA) followed by Tukey's multiple-comparison test; ns, not significant ($P \geq 0.05$); **, $P < 0.01$; ***, $P < 0.001$.

**Supplementary Table 1. Minimum inhibitory concentration of (*R*)-DI-87**

| Bacterial strain | MIC (µg/ml) |
|---|---|
| | **(*R*)-DI-87** |
| *S. aureus* Newman wild type | > 2048 |
| *S. aureus* Newman Δ*adsA* | > 2048 |

**Supplementary Table 2. Bacterial strains used in this study**

| Bacterial strain | Description | Reference |
|---|---|---|
| *E. coli* BL21 (DE3) pGEX-2T-*adsA* | BL21 bearing pGEX-2T-*adsA* expression plasmid | (Thammavongsa et al., 2009) |
| *S. aureus* Newman | Human clinical isolate | (Duthie and Lorenz, 1952) |
| *S. aureus* Newman Δ*adsA* | Newman Δ*adsA* | (Tantawy et al., 2022) |

**Supplementary Table 3. Cell lines used in this study**

| Cell line | Description | Reference |
|---|---|---|
| U937 | U937 cell line, ATCC® CRL-1593.2™ | ATCC |
| U937 $dCK^{-/-}$ | U937, bi-allelic deletion in *dCK* | (Winstel et al., 2018) |
